# Self-initiated restrictive feeding: a new mouse model of restrictive eating disorders

**DOI:** 10.64898/2026.10.20.760181

**Authors:** Robbie Ferguson, Mahesh Karnani

## Abstract

Restrictive eating disorders are serious psychiatric illnesses carrying the highest morality of any mental health condition, yet their neurobiology remains poorly understood. Progress is constrained by a shortage of animal models with convincing construct validity. In particular, anorexia nervosa (AN), a self-initiated restriction of caloric intake, is notoriously difficult to replicate in mice, which would otherwise offer an ideal pathway to understanding the neural mechanisms of this strongly gene-linked disorder, similar to advances in other neuropsychiatric conditions. Here, we introduce a model of self-initiated restrictive feeding (SIRF), using open-source behavioural devices. We show that SIRF specifically affects female mice, persists in the absence of the initial triggering conditions, and correlates with social rank – features that parallel AN. Meal pattern analysis shows that SIRF arises alongside decreased meal size and duration, with meal frequency unaltered in the long-term. Chemogenetic inhibition of the insular cortex, a brain region consistently implicated in AN and interoceptive processing, reduced SIRF. In sum, this proof-of-principle study suggests that SIRF provides a tractable model of self-initiated restriction as a platform for dissecting the neural circuits that initiate and sustain restrictive eating.

## Introduction

Restrictive eating disorders are a highly understudied and underfunded, yet urgent area of research (Beat, 2021; Deloitte Access Economics, 2020). Anorexia nervosa (AN) affects millions of people globally, with lifetime prevalence estimates of 1 - 4 % among women and 0.2 - 0.3 % among men, and has the highest mortality rate of all psychiatric disorders (Arcelus et al., 2011; Galmiche et al., 2019; van Eeden et al., 2021). While current mouse models of AN have questionable validity, the genetic tractability of mice and abundant neural circuit research tools present a promising pathway to understanding neural mechanisms of AN (Bulik et al., 2022). This is underscored by its increasingly well identified genetic risk loci (Grotzinger et al., 2026; Watson et al., 2019) and neural correlates (Foerde et al., 2022; Gaudio et al., 2018; Steinglass et al., 2019), and the relatively high (48 - 75 %) heritability and familial clustering of AN (Yilmaz et al., 2015). Furthermore, rodent models have enabled robust advances in other psychiatric disorders, like addiction and schizophrenia (Fuccillo et al., 2016). Previous rodent models of restrictive feeding have relied on involuntary food deprivation or mouse strains that eat less due to a genetic status (Méquinion et al., 2015; Scharner & Stengel, 2021) unrelated to identified polygenic risk alleles in AN (Watson et al., 2019). While these models can be useful in replicating aspects of starvation in ongoing AN, they cannot be used to investigate the biological mechanisms underpinning the initiation of AN.

A useful AN mouse model does not need to reproduce the full disorder, but should capture core features demonstrating several aspects of validity. The features we set out to reproduce were self-initiated restrictive feeding and a sex bias, and we define the relevant validity aspects as follows. *Face validity*: mice should, under ad libitum food availability, self-initiate sustained restriction of caloric intake despite preserved appetitive drive. *Aetiological validity*: restriction should be precipitated by environmental conditions with clinical analogues known to trigger AN, such as adolescent stress and exercise availability, rather than arising spontaneously or through imposed deprivation. *Construct validity*: restrictive feeding should show a sex bias consistent with human epidemiology, co-occur with objectively measurable correlates of AN such as social behaviour traits, and depend on brain regions and genes implicated in AN. *Predictive validity*: the model should respond appropriately to interventions with a known clinical signal and fail to respond to those shown ineffective in patients. Given the absence of established pharmacotherapy for AN, any model’s primary translational role should be to generate mechanistically grounded testable hypotheses for human trials. *Reliability*: testing conditions should be readily standardisable such that the phenotype can be reproduced across cohorts and sites.

Rodents in previous widely used AN models restore normal body weight after food access is restored (Dixon et al., 2003). Furthermore, while AN affects mainly women, previous mouse models have been more effective in males than females (Achamrah et al., 2017). As stressors play a central role in the development of eating disorders, and handling stress, e.g. during weighing and removal of feed, is a key factor reducing replicability of rodent studies, an ideal rodent model of AN would include handler-free weighing and food measurement. Another source of stress is single housing which is typical in previous AN models. As AN is associated with low sociability, perceived low social rank and submissive behaviours (Datta et al., 2021; Troop et al., 2003), measuring rodent social hierarchy is a useful addition to test model validity. We therefore set out to develop a behavioural paradigm that combines group-housing and several automated bio-behavioural measurements with plausible restrictive feeding triggering conditions to elicit Self-Initiated Restrictive Feeding (SIRF).

AN is typically triggered by stressful episodes during adolescence, which ultimately lead to a desire to control one’s body weight and food intake, or an outright fear of feeding (American Psychiatric Association, 2022). We induced such episodes by unpredictably delivering aversive air puffs to mice during feeding over three-day periods in the adolescent age range. This led to a robust transient decrease in feeding and body weight coupled with hallmarks of a fear of feeding, followed by the later development of decreased feeding and body weight, weeks after the air puff conditioning was terminated. We believe this model will be useful for further characterisation of brain areas involved in cognitive control of restrictive feeding. Among these regions, the insular cortex (IC) and prefrontal cortex (PFC) have repeatedly been implicated in AN through fMRI (Eddy et al., 2023; Foerde et al., 2015; Gaudio et al., 2018; Holsen et al., 2012; Uher et al., 2004), gross anatomy (G. K. Frank et al., 2013; G. K. W. Frank et al., 2016), and lesion cases (Uher & Treasure, 2005), and both regions have a monosynaptic pathway to the lateral hypothalamus (Razenberg et al., 2026), a well-established control centre of feeding (Kongstorp et al., 2025). As an initial step toward investigating these brain regions in our model, we chemogenetically inhibited the medial PFC (mPFC) or IC using the targeted expression of the hM4Di receptor (Roth, 2016) during air puff conditioning. IC inhibition led to a partial rescue of the long-term SIRF phenotype.

These results were made possible by a one-year grant from the Rosetrees Trust and the Stoneygate Trust. As funding for eating disorder research is scarce (Nimbley et al., 2026), we present these results at a very early stage in order to collaboratively develop this paradigm further. We therefore encourage feedback from patients, clinicians, researchers studying eating disorders, as well as those developing animal models, funders and potential industry partners.

## Methods

### Mice and husbandry

All experiments and procedures were approved by the University of Edinburgh local ethical review committee and performed under license from the UK Home Office in accordance with the Animal (Scientific Procedures) Act 1986. 48 wild-type C57BL6 mice were used in this study, aged 65-88 days at the time of the first induction epoch; data from Figure 2C to the end of the paper arose from age-matched groups 65-68 days old at the beginning of datasets. Mice were maintained on a reversed 12:12 hour light:dark cycle (lights off at 12:00 pm) and provided ad libitum access to food and water.

### Behavioural apparatus

For the duration of the experiment, mice were housed in an automated measurement enclosure (Figure 1A) in groups of 4 or 5 littermates. For all recordings of female mice, four identical enclosures were operated in parallel with age-matched mice.

**Figure 1.**
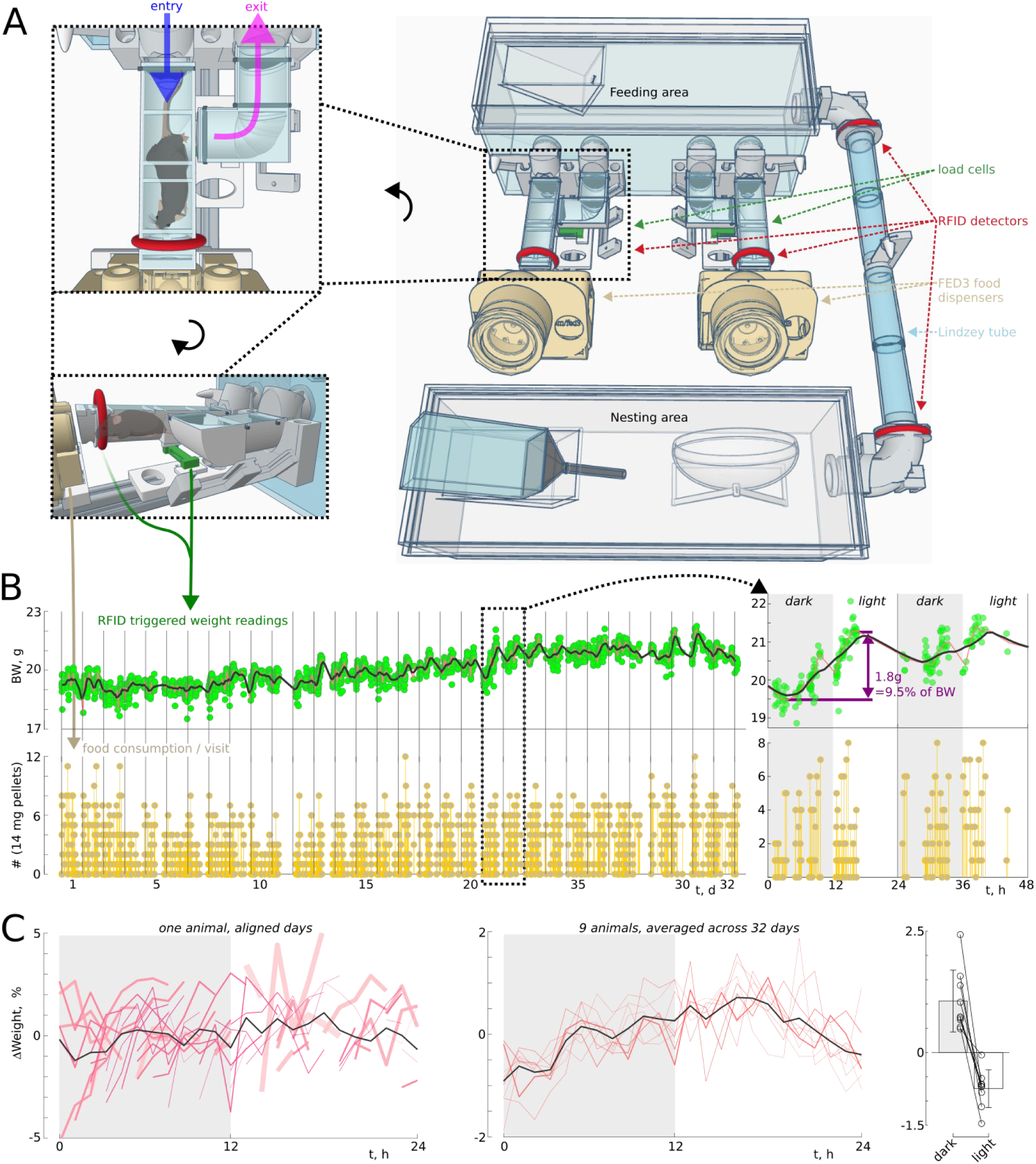
An automated device for handler-free measurement of body weight and food consumption. **A**, Schematic of the maze showing key functional features used to measure animal identity (RFID detectors), body weight (load cells), and social hierarchy (Lindzey tube), and to dispense food (FED3 food dispensers). **Top left inset** shows a top view of the feeding module with a mouse in the feeding tube, and **bottom left inset** shows the same module from a side angle showing the position of the load cell. The 3D model can be viewed at https://www.tinkercad.com/things/aLR9tcwqs0V-mazerex2. **B**, RFID triggered body weight (BW) readings and food consumption over 32 days in the enclosure from one female mouse aged 66 days on day 1. Inset shows expanded data over two days to highlight normal body weight fluctuations. **C**, **Left**, hourly average body weight change from daily median across each day plotted in different size and shade to highlight the typically observed large body weight changes during bouts of feeding, which could occur at any time of day. Average across days shown in black. **Middle**, averages across days for 9 animals (red), and the grand average across mice (black). **Right**, change in body weight during the dark and light epochs of each animal in the middle plot (last 3 h – first 3 h).

Each device consisted of a nesting enclosure: a standard 370*130*130 mm perspex cage with woodchip and paper bedding material, a running wheel and water. The nesting enclosure was connected through a ∼500 mm long 30 mm diameter tube made of end to end connected cut conical tubes and 3D-printed joinery. This feature forms the automated Lindzey tube which accumulated dominance hierarchy data throughout the experiment when spontaneous near-simultaneous entries of one mouse from each end occurred. To increase the likelihood of such occurrences, the ends of the tube contained radio-frequency identification (RFID) detectors, a beam-break sensor and one-way lockable doors which were operated to stop entry of multiple mice from one end. Thus when a mouse entered from each end, both mice typically exited from the same end in a sequence, with the last one being scored ‘winner’ of the interaction. In the data presented here (Figure 4), each mouse pair repeated these interactions 7.0 ± 3.7 times over the course of the recording. Social hierarchy was calculated from the full array of pairwise interactions using David’s score (Gammell et al., 2003) which takes into account the relative win rates of each individual.

At the other end of the Lindzey tube was the feeding enclosure: another standard perspex cage with only woodchip bedding. This cage had two feeding/weighing modules attached to it through 30 mm holes. These modules (Figure 1A inset) consisted of a feeding tube with an RFID detector, mounted on a 100 g load cell (Sparkfun TAL221) such that a weight reading could be taken when a mouse was detected. At the far end of the feeding tube, a modified feeding experimentation device 3 (FED3) pellet dispenser (Matikainen-Ankney et al., 2021) was placed where animals could retrieve 14 mg food pellets (Bio-Serv™ Dustless Precision Pellets™ for Rodents, F05684, 3.6 kcal/g) ad libitum. The FED3 was modified with a ‘cement mixer’ food hopper designed by Kravitz and colleagues, and a pressurised air conduit that guided a 30 PSI pressure line through the 3D-printed body to the front of the apparatus in order to deliver aversive non-harmful stimuli to the mice. 24 V coil valves were operated with a high-end switch and Arduino nano microcontroller to deliver 500 ms air puffs at 50 % likelihood during the induction epochs. To ensure that mice could not hide from the air puffs, we made it impossible to rapidly back out of the tube by unidirectional flaps at the entry of the feeding tube and within the curved exit tube leading back to the feeding area cage. The entry and exit were elevated 90 mm above the cage floor.

Data were recorded on a Raspberry Pi 5 computer, which interfaced with Sparkfun load cell amplifiers (NAU7802) and RFID detector amplifiers (SEN-15191) through a Qwiic Mux Breakout (TCA9548A), an Arduino nano board through USB, and a number of GPIO connections to the above boards and beam-break sensors (DFRobot SEN0503 and FC-51 type IR sensors). We will include a bill of materials, design files and build instructions for the technical community in the next version of this preprint, and will share more detail and materials upon request.

The device was restarted, de-dusted, restocked and calibrated once a day between 9 am and 12 pm. To keep this brief procedure minimally stressful, entry to the feeding tubes was prevented by sliding blockers.

### Surgery

All mice were injected intraperitoneally (i.p.) with a mixture of fentanyl (0.05 mg/kg), medetomidine (0.5 mg/kg) and midazolam (5 mg/kg), and an RFID tag (Sparkfun SEN-09416) was implanted under the chest skin using an injector implanter. For control and SIRF mice, anaesthesia was then antagonised with an i.p. injection of flumazenil (0.1 mg/ml) and atipamezole (5 mg/ml). For hM4Di mice, intracranial injections were performed before antagonising anaesthesia: mice were placed into a stereotactic apparatus and received a small subcutaneous drop of bupivacaine (0.25 %) under the scalp. The scalp was opened with one incision, the skull was levelled, and 0.5 mm craniotomies drilled. Using a needle edge, the deepest skull layer at each craniotomy site was removed to expose the surface of the brain. The injection needle was then placed on top of the brain and the depth co-ordinate set to zero. PFC-hM4Di animals were injected with 100 nl of 10^13^ vg/ml AAV9-hSyn-hM4Di-mCherry with flow rate 100 nl/min at three sites in each hemisphere: 2.2 mm anterior from Bregma, 0.4 mm lateral at a depth of 1.75 mm, and 1.8 mm anterior from Bregma, 0.4 mm lateral at depths 1.5 mm and 2.0 mm for the medial prefrontal cortex. IC-hM4Di mice received 100 nl injections at three sites in each hemisphere: 0.0 mm from Bregma, 3.8 mm lateral at a depth of 2.8 mm; 0.5 mm from Bregma, 3.5 mm lateral at depth 2.8 mm; and 1.0 mm from Bregma, 3.2 mm lateral at depth 2.7 mm. The injector was left in place for at least 2 minutes after each injection. The wound was stitched closed, and anaesthesia was antagonised. Mice were then left to recover for two weeks post-surgery in a home cage and then placed in the behavioural apparatus to habituate for one week before experimental interventions.

### Chemogenetic experiments

The hM4Di agonist C21 (Jendryka et al., 2019; Yu et al., 2024) was introduced into drinking water at 25 mg/L 15 h before the onset of each induction epoch in order to allow hM4Di activation to set in before induction. It was removed at the same time as induction ended.

### Histology

Viral targeting was assessed with post-experiment histological analysis. Mice received an i.p. overdose of pentobarbital, and were transcardially perfused with 4% paraformaldehyde. After 48 h post-fixation, 100 μm thick coronal sections were cut using a vibratome and mounted on glass slides. Slides were imaged using a 4x objective on a Zeiss Axio Scan.Z1 slides scanning microscope, and the extent of hM4Di-mCherry expression throughout the brain was determined by visually ascertaining presence of fluorescent somata in three equally spaced sections spanning the injected area, and tracing the expression pattern to coronal diagrams using QuPath software (Bankhead et al., 2017) (Figure 6A,B). All specimens had strong expression throughout the targeted areas: in mPFC-hM4Di mice infralimbic, prelimbic and anterior cingulate cortices; and in IC-hM4Di mice the insular, gustatory and visceral cortices. In addition, as shown in coronal diagrams in Figure 6A and B, weak off-target expression was detected in 3/9 mPFC-hM4Di mice in the medial septum, and in IC-hM4Di mice, there was typically some expression in adjacent cortical and subplate areas including the dorsal piriform cortex, ventral secondary somatosensory cortex, claustrum and endopiriform nucleus.

### Statistics

Data were analyzed using Matlab R2026a and reported as mean ± SD including in all bar graphs and errorbars. In Figure 5, meals were defined, based on analysis of inter-detection interval data and observations of how long an animal took to eat one food pellet (∼10 s), as RFID detections in the feeding tube separated in time by at least 30 s + 10 s for each pellet consumed. Meal size was computed as the number of 14 mg pellets consumed during each meal, and meal duration as the known duration between separate RFID detection during the meal if available + 10 s for each pellet consumed after the last RFID detection in a meal. At baseline, meals lasted 50.6 ± 12.2 s (n = 18 female mice), during which 4.9 ± 1.2 pellets, i.e., 247 ± 60 cal, were consumed, which is consistent with previous measurements with the same mouse strain and food dispenser (Barrett et al., 2025; Taghipourbibalan & McCutcheon, 2026).

For the chemogenetic experiments we used one-way repeated measures ANOVA with group (ctrl, SIRF, SIRF mPFC-hM4Di or SIRF IC-hM4Di) as the independent variable. When a significant group effect was found, three follow-up Student’s t-tests were used with Benjamini-Hochberg FDR corrected significance thresholds. All reported p-values are uncorrected such that readers can apply different corrections should they wish to, and all significance thresholds are Benjamini-Hochberg FDR corrected from 0.05. We used the following notation: ns = not significant, * = significant with p > 0.005, ** = significant with p < 0.005, significant with p < 10^-4^. As indicated, we performed statistical testing by one-way ANOVA, paired or two sample Student’s t-tests, Pearson correlations and linear regression fitted by ordinary least squares with coefficient significance assessment by two-sided t-tests.

## Results

### An open-source device for automated measurement of body-weight and food intake of group-housed mice

We modified existing open-source behavioural measurement devices (Hartmann et al., 2024; Matikainen-Ankney et al., 2021; Noorshams et al., 2017) to build a maze (Figure 1A) with separate feeding and nesting areas, achieving four features: 1) Individual mice are recognised by subcutaneous RFID tags which are detected with antennas in four locations. 2) In order to eat, mice must enter a tube that only fits one mouse at a time. This tube is on a load cell such that the weight of the mouse can be recorded several times a day. 3) FED3 food dispensers (Matikainen-Ankney et al., 2021) were adapted to contain a high-pressure line that can be controlled to deliver an aversive air puff when feeding is detected, and which cannot be evaded due to a one directional entry-exit system (Figure 1A, top left). 4) Mice have ample nesting materials, a running wheel and water in a separate enclosure which is accessed through a Lindzey tube which scores social hierarchy based on spontaneously occurring simultaneous entries from both ends of the tube. We used low-cost and open-source materials in order to democratise experimentation: 3D-printed parts (designs publicly available at https://www.tinkercad.com/things/aLR9tcwqs0V-mazerex2), the FED3, Raspberry Pi and Arduino microcontrollers, Sparkfun sensors, and a small number of other generic off-the-shelf components (see Methods).

Groups of 4-5 mice readily habituated to the maze, establishing nests in the nesting area and entering the feeding tubes ∼20-60 times each day to obtain all their food in the form of 14 mg complete rodent chow pellets (TestDiet precision pellet diet, 3.6 kcal/g). This resulted in a high density of body weight measurements that revealed marked weight fluctuations within each day of up to 9.5 % body weight, as well as normal growth trajectories over >1 month (Figure 1B). Mice tended to eat multiple meals of 247 ± 60 cal (n = 9 animals) in a span of a few hours such that their body weights increased in bouts which could happen at any time of the day but were more likely to occur during the dark phase (Figure 1C). This resulted in an average body weight increase of 1.1 ± 0.6 % (n = 9 animals, 32 days) during the dark phase and loss of 0.8 ± 0.4 % during the light phase (Figure 1C, Right). These results are consistent with previous high temporal resolution body weight monitoring results (Ahloy-Dallaire et al., 2019; Minematsu et al., 1991) demonstrating that our device is capable of measuring body weight and feeding without handler intervention.

### Induction of female-specific restrictive feeding using unpredictable aversive conditioning of feeding

After a minimum of 7 days habituation to establish a consistent baseline, we induced the SIRF phenotype by introducing unpredictable air puffs from the FED3 dispensers when a mouse withdrew a food pellet. Air puffs occurred at a maximally unpredictable 50 % likelihood and typically caused female mice to retreat from the feeding tube. On the first day of the 3-day induction period we observed female mice forming a huddle in the feeding or nesting areas, the frequency of detecting female mice in the feeding tubes dropped by 39 ± 55 % (n = 10 animals), and the amount of food consumed dropped by 81 ± 19 %. Male mice were not seen huddling, were detected at unaltered frequency in the feeding tubes (-4 ± 35 %, n = 10 animals) and their food consumption dropped only 15 ± 31 %. These marked behavioural changes during induction were reflected in a significant body weight loss in female but not male mice (Figure 2A,B). For the 3 days following the induction period, no further air puffs were administered, but female mice did not gain weight above their baseline weights, while male mice had gained significant weight (Figure 2A,B). Therefore, the SIRF phenotype is specific to female mice.

**Figure 2.**
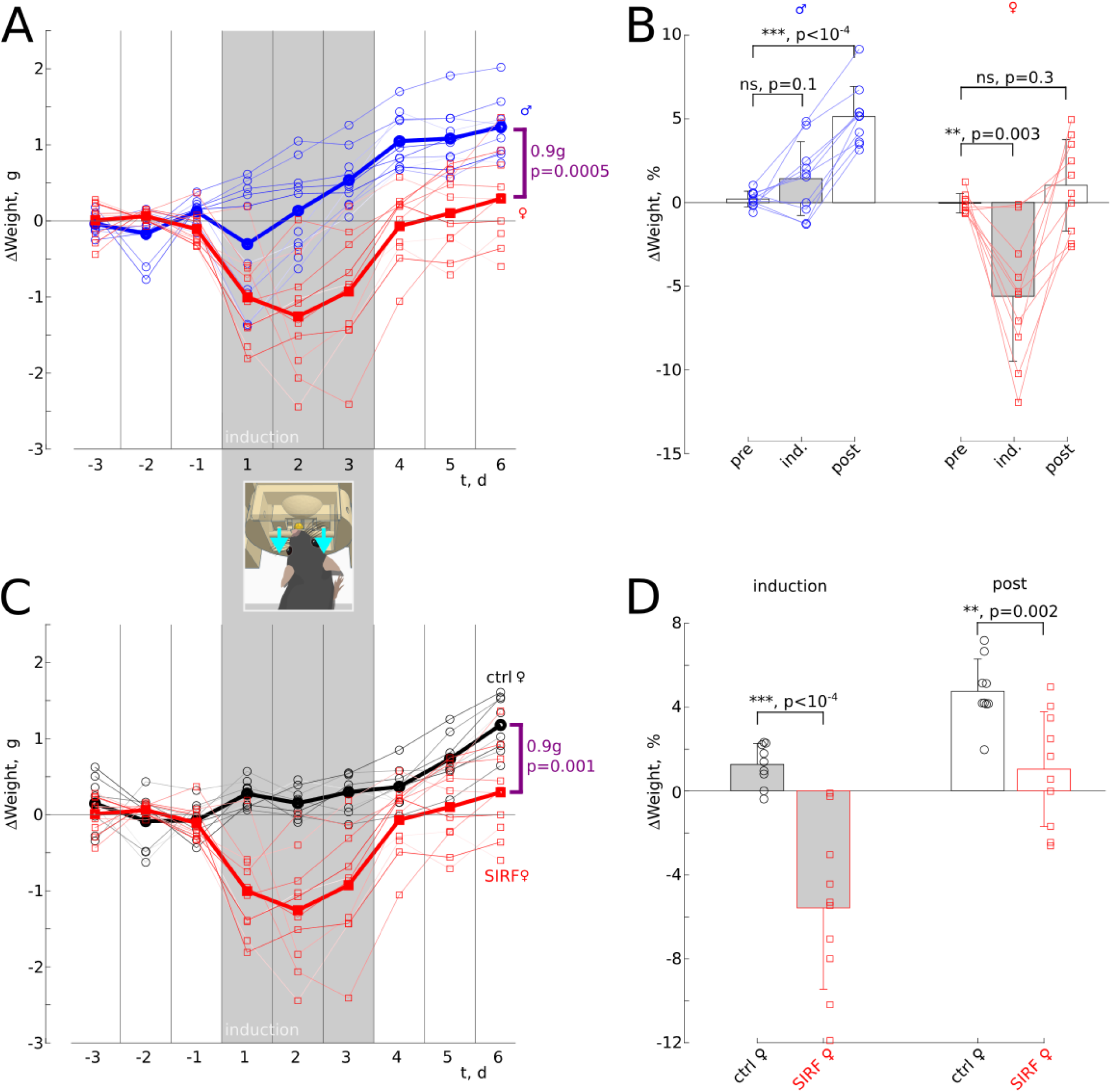
SIRF is female specific and out-lasts induction. **A**, Change in daily median weights of 10 male (blue, circles) and 10 female (red, squares) mice during 50 % likelihood facial air puff stimuli upon food retrieval during days 1-3 (induction, shaded, with cartoon illustrating in cyan arrows the air puffs from the modified FED3). Group mean plotted with a thick line. Difference between groups on last day indicated on plot (0.9 g, p = 0.0005 Student’s t-test). **B**, Average change in daily median weights from baseline in males (n = 10, blue circles) and females (n = 10, red squares) during two-day epochs before induction (pre, days -2 and -1), during it (ind., days 2 and 3), and after (post, days 5 and 6). Paired Student’s t-tests. **C**, As in A, for female mice subjected to induction (SIRF, red, squares, n = 10) and female controls not subjected to induction but measured in parallel in the same enclosures (ctrl, black, circles, n = 9). Difference between groups on last day indicated on plot (0.9 g, p = 0.001 Student’s t-test). **D**, Average change in daily median weights from baseline in ctrl females (n = 9, black circles) and SIRF females (n = 10, red squares) over two-day epochs during induction (induction, days 2 and 3), and after (post, days 5 and 6). Student’s t-tests.

We next compared female SIRF mice to age-matched control mice in the same measurement devices without exposure to air puffs. Compared to controls, SIRF mice lost body weight significantly during induction and remained below control body weight gain 3 days after induction (Figure 2C,D). This demonstrates that a lasting decrease in body weight gain was triggered by the induction protocol in female SIRF mice. Notably, both induction and maintenance of effects on body weight were in the presence of continuous access to food.

### Persistent SIRF develops weeks after induction

AN is a chronic condition, and repeated stressful episodes are a significant predictor of symptom provocation (Berry et al., 2024; Grilo et al., 2012). Therefore, we repeated the induction period, and monitored female mice over longer time courses (Figure 3). Body weight gain trajectories of control and SIRF groups diverged from day 17 after the onset of the first induction epoch, and by the end of the 25-day experiment SIRF mice had gained significantly less weight (1.6 ± 0.8 g, n = 10) than control mice (2.4 ± 0.7 g, n = 9, p = 0.03 Student’s t-test; Figure 3A). As we noticed this late emerging difference after the first of two experimental runs, we monitored the second experimental run for an additional week, and observed a continued divergence in weight gain (at end of experiment: SIRF 1.3 ± 0.8 g, n = 5; ctrl 2.6 ± 0.8 g, n = 4, p = 0.03 Student’s t-test; Figure 3B). At the the end of the 25-day experiment, this was accompanied by decreased daily food consumption of 17 ± 24 % in SIRF animals (n = 10) and an increased daily food consumption of 15 ± 14 % in controls (n = 9, p = 0.003 Student’s t-test, Figure 3C). In the animals recorded for an additional week, this difference persisted (at end of assay: SIRF -30 ± 13 %, n = 5; ctrl 4.2 ± 14 %, n = 4, p = 0.007 Student’s t-test; Figure 3D).

**Figure 3.**
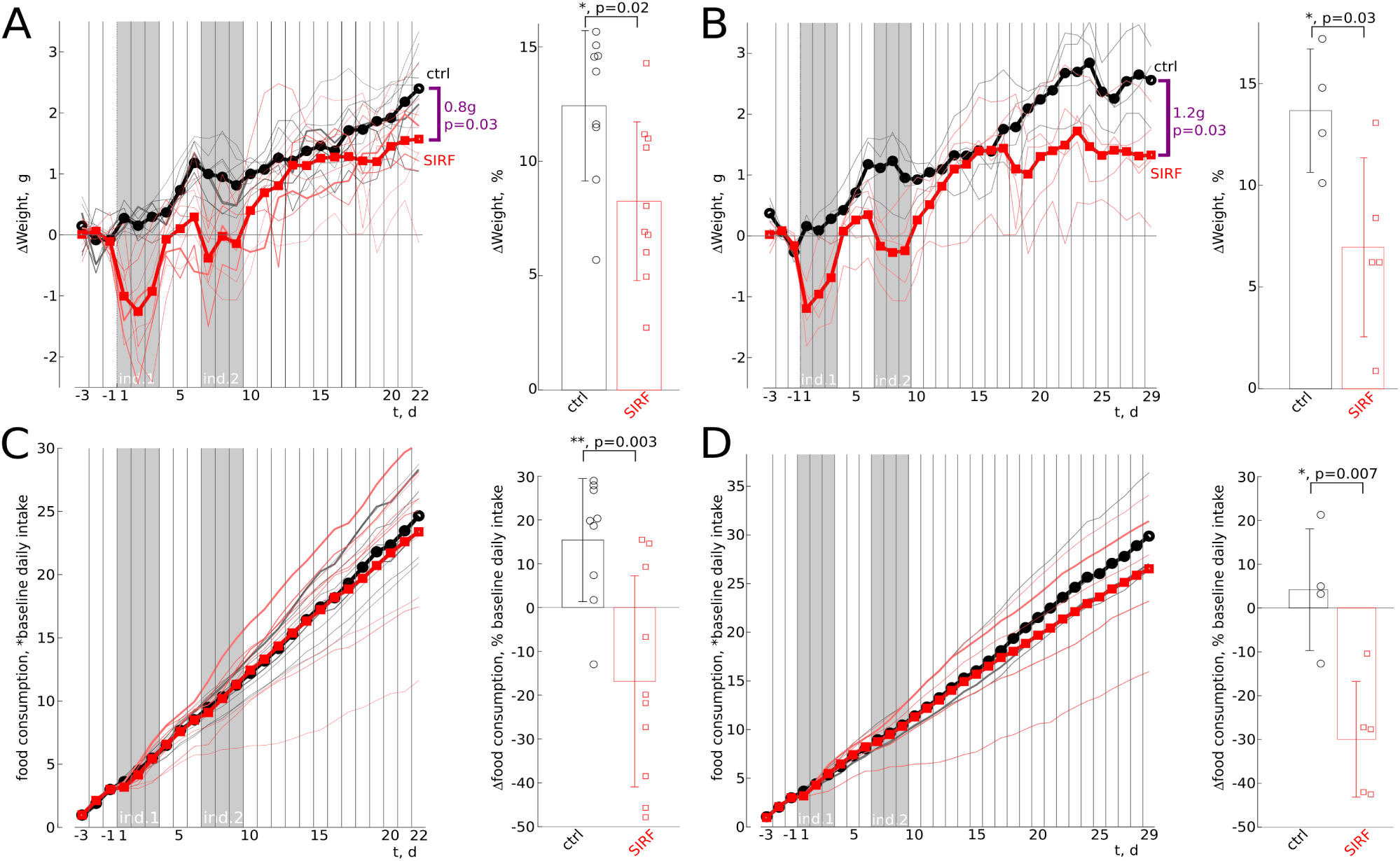
long-term SIRF. **A**, **Left**, Change in daily median weights from baseline in ctrl females (n = 9, black circles) and SIRF females (n = 10, red squares) during and after 50 % likelihood facial airpuff stimuli upon food retrieval during days 1-3 and 7-9 (ind. 1 and ind. 2, shaded). **Right**, % change from baseline by two last days of the dataset (days 21 and 22). **B**, As in A, but for a subset of mice that were recorded for an additional week (ctrl n = 4, SIRF n = 5). **Right**, % change from baseline by two last days of the dataset (days 28 and 29). **C, Left**, Cumulative food consumption as fraction of baseline (days -3 to -1) daily intake for same animals as in A. **Right**, % change from baseline over two last days of the dataset (days 21 and 22). **D, Left**, As in C but for mice in B. **Right**, % change from baseline over two last days of the dataset (days 28 and 29). All comparisons by Student’s t-tests.

### SIRF correlates with social rank

AN patients report lower perceived social status, negative opinions of friendships and more self-reliance than healthy control subjects (Datta et al., 2021; Troop et al., 2003). In mice, such perceived social dimensions cannot be measured, but social hierarchy can be measured objectively using the Lindzey tube test (van den Berg et al., 2015). We found that social rank correlates with the weight gain in SIRF mice (Spearman correlation 0.64, p = 0.046, n = 10; Figure 4). However, the correlation appears inverse from the self-reported submissive traits of AN patients, in that socially dominant mice had more severe SIRF phenotypes. This may be explained through the difference in self-reported and objective measures of social status (see Discussion).

**Figure 4.**
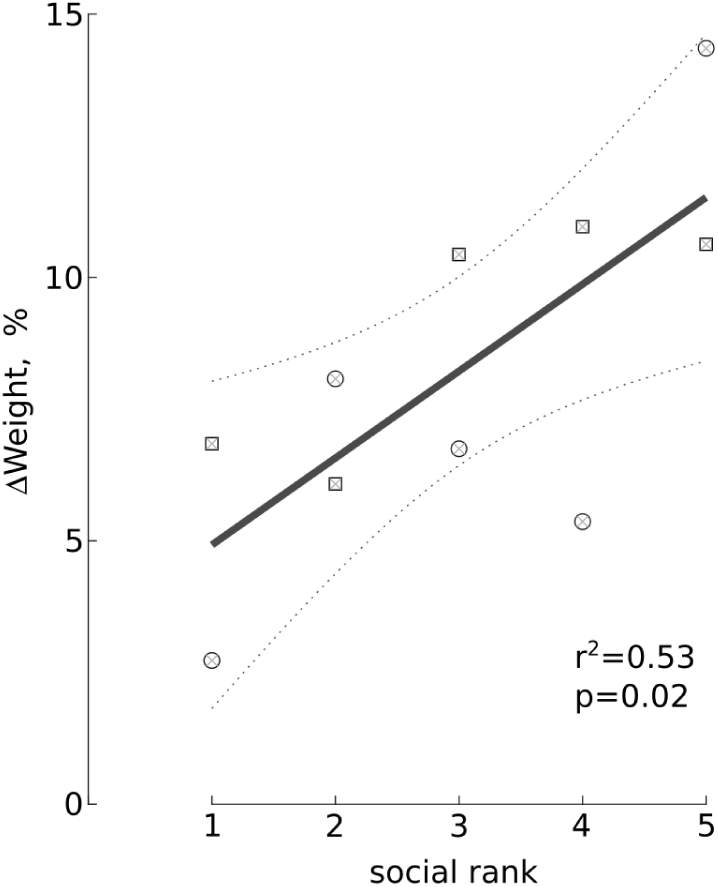
socially dominant female mice are more susceptible to long-term SIRF. Social hierarchy rank (1 = most dominant, 5 = most submissive) as measured by repeated spontaneous Lindzey tube tests in the enclosure and quantified as David’s score in the two groups (circles and squares) of 5 female SIRF mice and their weight changes from Figure 3A (Right panel). Linear fit (heavy line) and 95% confidence bounds (dotted lines). Linear regression was fitted by ordinary least squares and coefficient significance assessed by a two-sided t-test.

### Meal structure alterations during SIRF

To gain further insight into the factors driving SIRF, we analysed meal structure throughout the experiment (Figure 5). SIRF and control females had indistinguishable meal sizes at baseline (ctrl 4.9 ± 0.9 pellets, n = 9; SIRF 4.8 ± 1.4 pellets, n = 10; p = 0.9 Student’s t-test), which decreased in SIRF mice during induction (ctrl 4.9 ± 0.8 pellets; SIRF 1.9 ± 0.5 pellets; p < 10^-7^ Student’s t-test) and remained low compared to controls (at end of assay, ctrl 5.7 ± 1.3 pellets; SIRF 4.5 ± 0.6 pellets; p = 0.01 Student’s t-test, Figure 5A). Similarly, meal duration was initially indistinguishable between the groups (ctrl 52 ± 11 s; SIRF 49 ± 15 s; p = 0.7 Student’s t-test), decreased during induction in SIRF mice (ctrl 50 ± 8.0 s; SIRF 28 ± 9.6 s; p < 10^-4^ Student’s t-test), and remained low (ctrl 59 ± 12 s; SIRF 44 ± 7.2 s; p = 0.005 Student’s t-test, Figure 5B). These data suggest that SIRF mice are cautious eaters, and their shorter meals may underlie their diminished weight gain.

**Figure 5.**
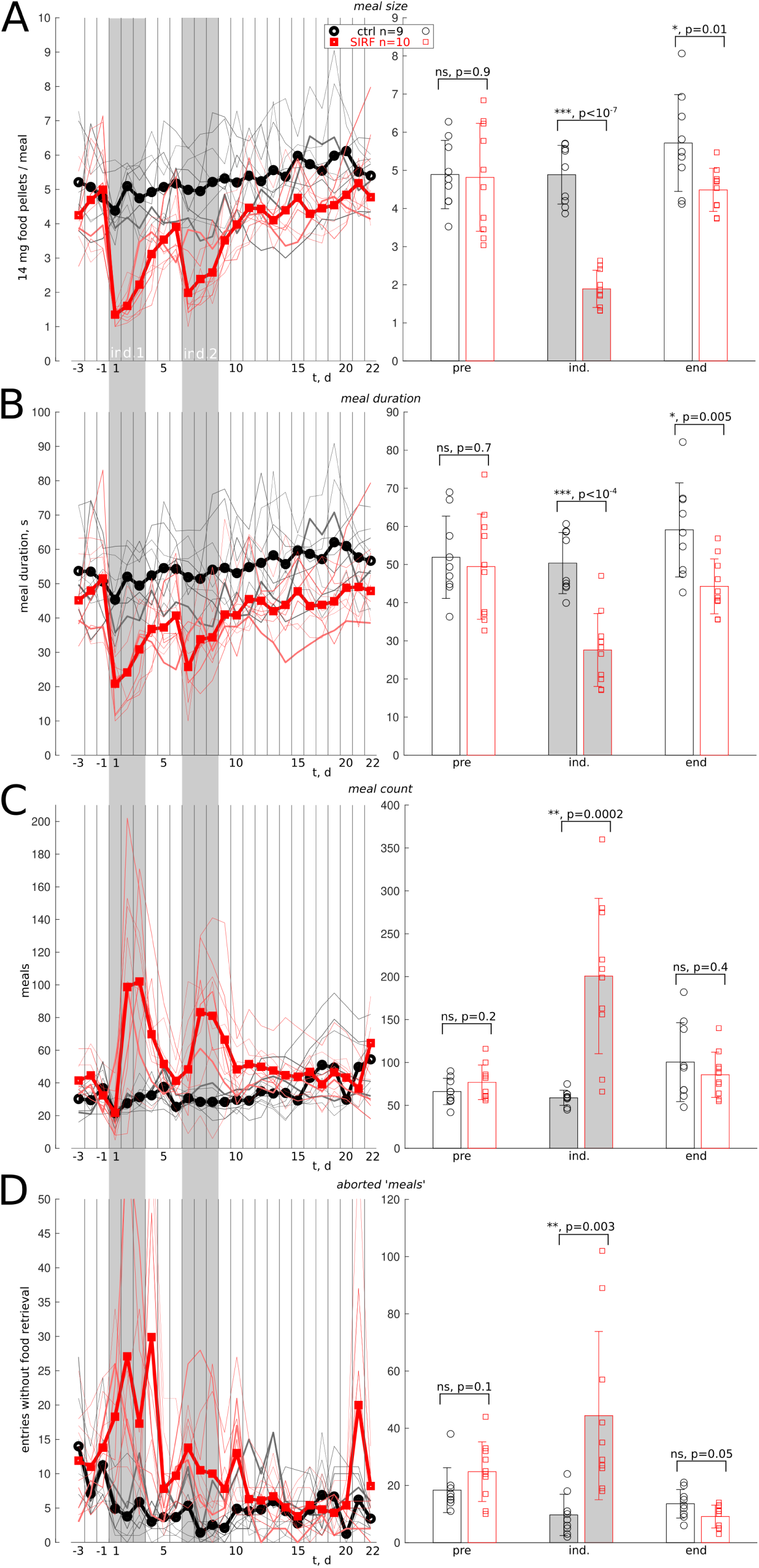
meal structure changes during SIRF. **A, Left**, Mean meal size (amount of 14 mg food pellets) in control females (n = 9, black lines and circles) and SIRF females (n = 10, red lines and squares). **Right**, mean meal size from continuous two-day epochs before induction (pre, days -2 and -1), during it (ind., days 2 and 3), and end of assay (end, days 18 and 19). **B**, As in A, but mean meal duration. **C**, as in A, but number of meals (see Methods). **D**, as in A, but aborted ‘meals’, i.e., entries to the feeding tube that would be classified as separate meals in terms of time from temporally adjacent entries but during which no food was retrieved. All comparisons by Student’s t-tests.

As noted above, the number of entries into the feeding tubes were initially decreased on the first day of SIRF induction, and this was visible also in the number of meals (Figure 5C). However, as induction continued, meal counts increased robustly, but were similar between control and SIRF mice at the end of the experiment (Figure 5C). As not all entries into the feeding tubes were meals, and entries without feeding can be interpreted as a fear of feeding, we quantified the number of such aborted ‘meals’ (Figure 5D). These were similar in baseline (ctrl 18 ± 7.8; SIRF 25 ± 10; p = 0.1 Student’s t-test), increased in SIRF mice during induction (ctrl 9.7 ± 7.2; SIRF 44 ± 29; p = 0.003 Student’s t-test), and not significantly different in the long-term (ctrl 14 ± 5.0; SIRF 9.1 ± 4.0; p = 0.05 Student’s t-test). Taken together, these results suggest that SIRF mice terminate their meals earlier than controls and may have a fear of feeding that drives their diminished weight gain, a condition with strong face validity for AN.

### Inhibition of cortical regions implicated in restrictive eating disorders

Given that SIRF exhibits strong face validity as a model of AN, we sought to test if manipulating brain regions implicated in AN would impact the development of SIRF. As there is little information on which brain regions could underlie initiation of AN, we sought a loss-of-function in brain regions known to show differences in ongoing AN and recovered patients. To do this, we used stereotactic injections of AAV9-hSyn-hM4D(Gi)-mCherry to drive expression of the chemogenetic neuronal inhibitor hM4Di (Roth, 2016) in the medial prefrontal cortex (SIRF mPFC-hM4Di group, Figure 6A) or insular cortex (SIRF IC-hM4Di group, Figure 6B). These cortices have consistently been implicated in AN (Eddy et al., 2023; Gaudio et al., 2018; Holsen et al., 2012, 2014; Uher et al., 2004; Uher & Treasure, 2005), and project functional glutamatergic synapses to the lateral hypothalamus (Razenberg et al., 2026), a well-known controller of food consumption. To uninvasively inhibit the targeted cortical regions during the 3-day induction epochs, we supplemented drinking water with the hM4Di agonist C21 (25 mg/l) which has been shown to be an effective administration route (Yu et al., 2024).

**Figure 6.**
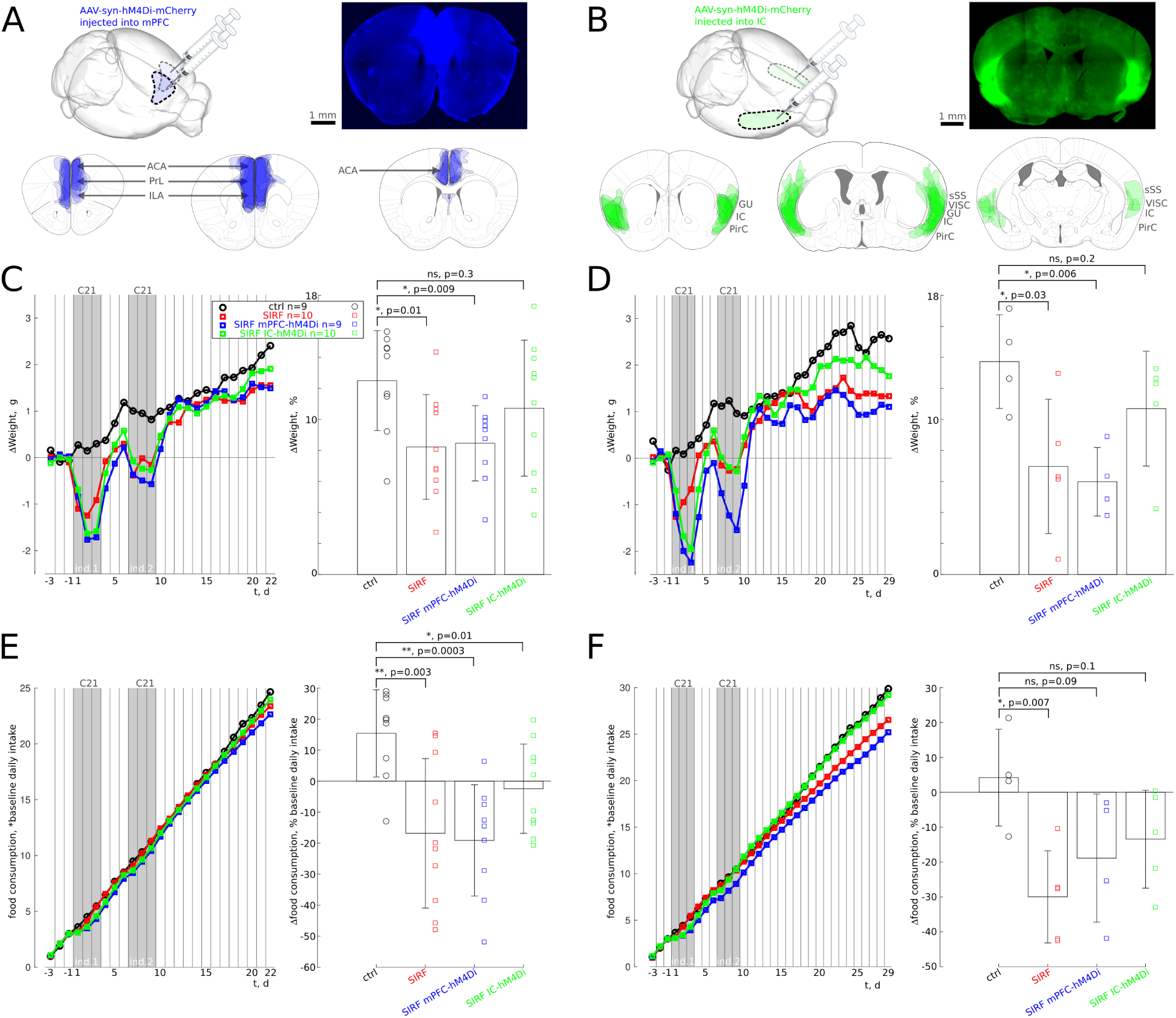
Chemogenetic inhibition of cortical targets of AN. **A, Top left**, Schematic of AAV-syn-hM4Di-mCherry injection into mPFC. **Top right**, example coronal section from the middle antero-posterior (AP) coordinate of the injection site, mCherry fluorescence pseudocoloured blue. **Bottom**, tracings of the extent of expression in all animals (n = 9) at AP levels (from Bregma), +2.2 mm, +1.4 mm, and +0.7 mm. **B, Top left**, Schematic of AAV-syn-hM4Di-mCherry injection into IC. **Top right**, example coronal section from the middle antero-posterior (AP) coordinate of the injection site, mCherry fluorescence pseudocoloured green. **Bottom**, tracings of the extent of expression in all animals (n = 10) at AP levels (from Bregma), +1.2 mm, -0.1 mm, and -1.1 mm. **C**, **Left**, Change in daily median weights from baseline in ctrl females (n = 9, black circles) and SIRF females (n = 10, red squares) SIRF mPFC-hM4Di females (n = 9, blue squares) and SIRF IC-hM4Di females (n = 10, green squares) during and after combined SIRF induction (not controls) and chemogenetic agonist (C21 via drinking water to all groups including controls) administration during days 1-3 and 7-9 (ind. 1 and ind. 2, shaded). **Right**, % change from baseline by two last days of the dataset (days 21 and 22). **D**, As in A, but for a subset of mice that were recorded for an additional week (ctrl n = 4, SIRF n = 5, SIRF mPFC-hM4Di n = 4, SIRF IC-hM4Di n = 5). **Right**, % change from baseline by two last days of the dataset (days 28 and 29). E**, Left**, Cumulative food consumption as fraction of baseline (days -3 to -1) daily intake for same animals as in A. **Right**, % change from baseline over two last days of the dataset (days 21 and 22). F**, Left**, As in C but for mice in B. **Right**, % change from baseline over two last days of the dataset (days 28 and 29). All comparisons by Student’s t-tests after a one-way ANOVA omnibus test across all groups with p < 0.05.

Each SIRF group responded similarly to the induction protocol, with decreased body weight and feeding (Figure 6C). A one-way ANOVA test revealed a significant effect of group on body weight gain by end of experiment (F(3, 34) = 3.19, p = 0.04, Figure 6C), and post-hoc Student’s t-tests revealed that only the SIRF and SIRF mPFC-hM4Di had significantly lower body weight gain than controls, by approximately 30 % (ctrl 12 ± 3.2 %, n = 9; SIRF 8.2 ± 3.4 %, n = 10; SIRF mPFC-hM4Di 8.5 ± 2.4 %, n = 9; p < 0.01), whereas body weight gain of the SIRF IC-hM4Di group was not significantly different from control (11 ± 4.4 %, n = 10; p = 0.3). To test for longer term effects, we again tested separately the second experimental run that was recorded for an additional week, and found the same result (one-way ANOVA F(3, 14) = 4.33, p = 0.02 and post-hoc Student’s t-tests as in Figure 6D).

All SIRF groups had significantly lower increases in food intake than controls, although SIRF IC-hM4Di mice showed the lowest effect at nearly 50 % of the effect on SIRF and SIRF mPFC-hM4Di mice (one-way ANOVA F(3, 34) = 7.03, p = 0.0008; ctrl 15 ± 14 %, n = 9; SIRF -17 ± 24 %, n = 10; SIRF mPFC-hM4Di -19 ± 18 %, n = 9; SIRF IC-hM4Di -2.5 ± 14 %, n = 10; all p < 0.01 compared to ctrl, Figure 6E). In the animals recorded for an additional week, an overall group effect persisted (one-way ANOVA F(3, 14) = 4.04, p = 0.03), however only SIRF mice decreased food consumption significantly compared to controls (ctrl 4.2 ± 14 %, n = 4; SIRF -30 ± 13 %, n = 5; p = 0.007 Student’s t-test; Figure 6D), while both cortical loss-of-function groups were not significantly different from control (SIRF mPFC-hM4Di -19 ± 18 %, n = 4; SIRF IC-hM4Di -13 ± 14 %, n = 5; p > 0.09 compared to ctrl by Student’s t-test; Figure 6F). Taken together, these results suggest that inhibiting the IC during SIRF induction partially rescues the SIRF phenotype by normalising body weight gain, and limited impact on the reduction in food intake. Given that the insular cortex is among the most consistently altered brain structures in AN (Holsen et al., 2012, 2014; Uher et al., 2004), and one of the few structures shown to be altered in the early stages of AN (Gaudio et al., 2018), this finding supports the construct validity of SIRF as an AN model.

## Discussion

Here we set up a new mouse model of restrictive eating disorders, SIRF. The method is based on fearful stimuli, which are often seen as a valid feature of rodent models of AN (Ulrich-Lai et al., 2015). However, a rodent model utilising a fear of feeding has not been established as a standard approach (Méquinion et al., 2015; Scharner & Stengel, 2021). Therefore, we summarise the distinctive aspects of SIRF which to our knowledge have not been used in previous studies with aversive stimulus suppression of feeding (Bodnoff et al., 1988; Mongeau et al., 2003; Petrovich & Lougee, 2011; Reppucci et al., 2013; Reppucci & Petrovich, 2018): 1) The nature of the aversive stimuli – we used maximally unpredictable (50 % likelihood) air puff stimuli delivered over three consecutive days, while previous studies have used 100 % likelihood foot shocks over a few training sessions to form a fear-inducing conditioned stimulus. Air puffs are similarly non-noxious, aversive, and as stressful as electrical shocks, but are also seen as a welfare improvement as they can be less painful can be escaped in principle (Lamy et al., 2016). Thus, the aversive stimulation in SIRF couples feeding with a state of fear and stress over a painless, yet uncertain and aversive outcome. We invite commentary on whether this approximates conditions triggering AN, and how to improve the model. 2) Continuous, handler-free monitoring – while previous AN rodent models have used manual handling to weigh animals, move them in and out of testing chambers, and remove and replace food, we used an increasingly common approach of building an extended home-cage device that automates measurement based on RFID tags (Bolaños et al., 2017; Erskine et al., 2019; Hartmann et al., 2024; Kiryk et al., 2020; Noorshams et al., 2017; Puścian et al., 2016; Puścian & Knapska, 2022; Silasi et al., 2018; Torquet et al., 2018; Woodard et al., 2017, 2020). This approach eliminates both handling stress and the discontinuity and potential variability inherent in test-session-based approaches. SIRF appears to be a promising model of restrictive eating disorders in general, and AN in particular, given the sex bias on female mice (Figure 2A), the long-term impact on food intake and body-weight gain (Figure 3), which correlated with social dominance (Figure 4). In addition, meal structure was altered in a way consistent with a fear of feeding (Figure 5), and inhibiting the insular cortex partially rescued long-term SIRF (Figure 6). These aspects contribute to face, construct and aetiological validity of SIRF as a preclinical model of the restricting subtype of AN. Additionally, given that SIRF relies on tactile aversive stimuli (air puffs) during feeding, it has face validity for the more recently characterised, and less well understood, avoidant/restrictive food intake disorder (ARFID), where aversive tactile sensation during feeding is a typical symptom.

Meal structure data are seldom reported in restrictive feeding models, and therefore our device and preliminary findings (Figure 5) break new ground – however, to the best of our knowledge, meal structure analyses in human AN patients is also scarce. The two studies we found show that AN patients over-estimate the size of portions (Milos et al., 2013) and eat larger meals over longer time course than healthy controls (Sunday & Halmi, 1996). The latter is a perplexing finding which may be due to the monitoring conditions, however, AN patients also had larger variability in meal size than healthy controls. Overall, our results suggest that SIRF mice terminate meals early, potentially due to mounting fear of negative consequences of eating. Further human studies could help elucidate if this enhances the aetiological validity of SIRF.

AN is associated with impairments across domains within social functioning and social cognition (Schmidt et al., 2025). Some of this evidence points to a correlation with social submissiveness. On one hand, AN patients report lower perceived social status, negative opinions of friendships and more self-reliance than healthy control subjects (Datta et al., 2021; Troop et al., 2003), suggesting submissive traits. One the other hand, a longitudinal study linking school age social status to likelihood of AN symptoms later in life, showed elevated AN likelihood in both bullies, victims and victim-bullies (Copeland et al., 2015), suggesting that also dominant and sub-dominant traits correlate with AN. Self-reported social attitudes should be tempered by the known low correlation between objective and subjective social status (Evans & Kelley, 2004; Richter & Menold, 2024), and complications arising from multiple reference points across various measures (Redhead & Power, 2022). We found a significant negative correlation between the severity of long-term SIRF and social dominance (Figure 4); more dominant mice were more susceptible to SIRF. This may align with over-controlling personality styles in AN (Isaksson et al., 2021), but, so far and to the best of our knowledge, evidence for objective human dominance hierarchy position (analogous to that in mice) correlating with AN symptoms is lacking.

The neural mechanisms underlying initiation of AN are a critical unknown (Steinglass et al., 2019), where rodent models provide critical experimental access. We have taken the first step here through cortical loss-of-function during SIRF induction, which points toward a role for IC and not mPFC. Taste activation of IC is lower in recovered AN patients (Oberndorfer et al., 2013; Wagner et al., 2008), along with altered interoceptive task activity (Kerr et al., 2016), pain processing (Strigo et al., 2013) and higher reward omission responses in active AN patients (G. K. W. Frank et al., 2012). A key study described decreased connectivity in a network involving IC in early stage AN (Gaudio et al., 2018), suggesting IC may be involved in the initiation of illness. Our preliminary findings that reduced activity in IC during the induction of SIRF decreases it, are consistent with an overall model of AN initiation through a process occurring within the IC, or driven by it in other structures, such as through insular projections to the hypothalamus (Razenberg et al., 2026). Additionally, altered insular activation has been found in ARFID (Kerem et al., 2022). On the other hand, mPFC may instead have more to do with maintenance rather than initiation mechanisms. We aim to dissect this further in future studies aimed at manipulation of these and other key brain structures (Foerde et al., 2015, 2022; Lloyd et al., 2023) during both initiation and maintenance stages of SIRF.

## Acknowledgements

This research would not have been possible without generous funding from the Rosetrees Trust and the Stoneygate Trust (Seedcorn2024\100100). We are very grateful for discussions that shaped this project with Drs Karin Foerde, Joanna Steinglass, Tim Walsh, Karri Gillespie-Smith, Helen Sharpe, Emy Nimbley, Fiona Duffy and Patrick Spooner. The original idea for the unpredictable airpuff was from Dr Rebecca Jordan. We thank Andre Borba, Vinicius Borges, Martyna Stasiak, Chris Wilson, Ryan Krones, Sara Wallace, Daniel Vanbeck Smith and Jamie Kean for expert technical assistance testing previous prototype versions related to this project.

